# Is *Drosophila melanogaster* Stress Odorant (dSO) really an alarm pheromone?

**DOI:** 10.1101/534719

**Authors:** Ryley T. Yost, Emerald Liang, Megan P. Stewart, Selwyn Chui, Andrew F. Greco, Shirley Q. Long, Ian S. McDonald, Jeremy N. McNeil, Anne F. Simon

## Abstract

Social interactions are crucial for the reproduction and survival of many organisms, including those using visual, auditory and olfactory cues to signal the presence of danger. *Drosophila melanogaster* emits an olfactory alarm signal, termed the *Drosophila* stress odorant (dSO) in response to mechanical agitation or electric shock, and conspecifics avoid areas previously occupied by stressed individuals. However, the contextual, genetic and neural underpinnings of the emission of, and response to dSO, have received little attention. Using a binary choice assay, we determined that neither age and sex of emitters, nor the time of the day, affected the emission or avoidance of dSO. However, both sex and mating status affected the response to dSO. We also demonstrated that dSO was not species specific so it should not be considered a pheromone but a general alarm signal for Drosophila. However, the response levels to both intra and inter-specific cues differed between species and possible reasons for these differences are discussed.

**HIGHLIGHTS:**

- Emission of dSO, a highly volatile chemical blend emitted by stressed flies, is not context specific
- Response to dSO is context specific, affected by factors such as age and mating status.
- As flies respond to volatiles for stressed heterospecifics, dSO should not be considered an alarm pheromone, but as an alarm cue.

## INTRODUCTION

Social interactions using visual, auditory, olfactory and/or tactile cues are crucial for the successful development, survival and reproduction of organisms (Dahanukar & Ray, 2011; Sokolowski, 2010), including those relating to danger. Reaction to dangers can elicit a range of different behavioural responses (Yew & Chung, 2015), involving different sensory modalities, which depends on the species and the ecological context (Verheggen, Haubruge, & Mescher, 2010). Olfactory alarm signals are typically made up of highly volatile, non-persistent molecules (single compounds or as a blend), which rapidly inform conspecifics of potential danger without generating a persistent state of alert (Verheggen et al., 2010). Alarm pheromones, which by definition modulate interactions between conspecifics, have been reported in a wide range of animals, from nematodes to humans (Chao, Fleischer, & Yang, 2018; Hunt, 2007; Mathuru et al., 2012; Mujica-Parodi et al., 2009; Vandermoten, Mescher, Francis, Haubruge, & Verheggen, 2012; Zhou et al., 2017), although in some cases these olfactory cues elicit responses in closely related sympatric species sharing common natural enemies (Napper & Pickett, 2008).

In *Drosophila melanogaster* stressed individuals emit an olfactory alarm cue, the *Drosophila* stress odorant (dSO) and flies avoid areas previously occupied by stressed conspecifics (Suh et al., 2004). Carbon dioxide (CO_2_) is one component of dSO, and the sensory pathway for the perception of CO_2_ has been investigated (Bracker et al., 2013; Faucher, Forstreuter, Hilker, & de Bruyne, 2006; Krause Pham & Ray, 2015; Kwon, Dahanukar, Weiss, & Carlson, 2007; Siju, Bracker, & Grunwald Kadow, 2014; Suh et al., 2007; Suh et al., 2004; Turner & Ray, 2009). However, the avoidance response is weaker to CO_2_ alone than to dSO suggesting other components are present (Suh et al., 2004). We know that both sexes respond to dSO, although the levels of response vary as a function of the genetic background (Fernandez et al., 2014) and age (Brenman-Suttner et al., 2018), as well as the density of emitters (Fernandez et al., 2014). In order to facilitate the identification of the complete blend of dSO, as well as the genes and sensory pathways involved, it is essential to know the conditions that result in high production of the olfactory cue. Therefore, we conducted experiments examining the effect of different factors on the emission and perception of dSO.

## METHODS

### Experimental animals

All *D. melanogaster* (Canton-S) and *D. simulans and D. suzukii* adults used in the different assays were obtained from laboratory colonies maintained at Western University. *D. melanogaster* and *D. simulans* were reared on Drosophila Jazz-Mix™ diet (Fisher Scientific, ON, Canada) at 25°C, 50% RH under a 12L:12D light cycle. *D. suzukii* was maintained on a banana-cornmeal-agar medium (Jakobs, Ahmadi, Houben, Gariepy, & Sinclair, 2017) at 21°C, 65% RH under a 14L:10hD light cycle. Virgins were obtained by sexing flies at emergence and holding adults in same sex containers (approximately 40 flies/container) until needed. To obtain mated flies, newly emerged flies (20 male and 20 females) were held together until tested.

### dSO Avoidance Assays

Unless otherwise stated, all experiments used mated adults between 3-7 days old, and were carried out at 25°C and 50% RH between 12:00 and 16:00 to reduce variation associated with any diel periodicity in behaviour (Dubruille & Emery, 2008). A detailed description of the dSO avoidance assay can be found in (Fernandez et al., 2014). In short, responder flies are placed in a binary-choice T-maze under uniform light conditions where they have a choice between a control vial and either one containing dSO produced by stressed flies that had been vortexed over 1 minute (15 seconds on, 5 seconds at rest, repeated 3 times) or a vial previously occupied by non-stressed flies. After 1 min we recorded the position of all flies and then calculated the performance index (PI) by subtracting the number of responder flies in the experimental vial from the number of flies in the air vial, divided by the total number of flies used in the assay and multiplied by 100. This provides a measure of the preference, where a positive value indicates responders avoided the experimental vial while a negative value indicated they were attracted to it. Thus, total avoidance of the experiment vial would give a PI of 100, while if flies were equally divided between the control and experimental vials (no preference) the PI would be 0.

When cues from undisturbed flies were required we used the counter-current apparatus first reported by Benzer (1967), as described in Fernandez *et al.* (2014). Flies were drawn from a shaded holder vial into a lit clean vial using a 15W white light, where they remained for 1 min, before being removed using the same technique.

Previous experiments (Fernandez et al., 2014) found that 3-7 day old mated flies avoided dSO produced by either sex alone or with a mixed sex population. However, it was unknown if mating status affected responses so we compared the responses of mated and virgin flies using 30 mated or virgin responders (all male, all female or 15 males and 15 females) to a dSO source produced by 70 emitters (single or mixed sexes). These were the same densities tested in previous studies performance but as the index decreases with the number of emitters (Fernandez et al., 2014), we repeated the experiment using 20 emitters and 15 responders.

To test the effect of age on dSO emission we carried out 9 replicates where 30 flies (mixed sex) were exposed to the cues from 70 adults, ranging in age from 2-5 days up to 7 weeks old. We also tested the sex of emitters by exposing 15 mixed sex responders to dSO obtained from either 20 males, 20 females or a mix of both, with a minimum of 9 replicates for each combination. We investigated the influence of responder age by exposing 15 responders to the dSO obtained from 20 males or females that were either 3-4 or 7-10 days old. There were 9 replicates per treatment.

To determine if there might be a diel periodicity in either the emission of or response to dSO we maintained two colonies under a 12L:12D photoperiodic regime, but with the lights-on signal at 04:00 in one and at 08:00 in the other. This allowed us to test different combinations for both emitters and responders, as shown in **Figure 1**, where ZT refers to the time of the lights on signal. There were 9 replicates for all combinations, with sexes being tested separately.

**Figure 1.**
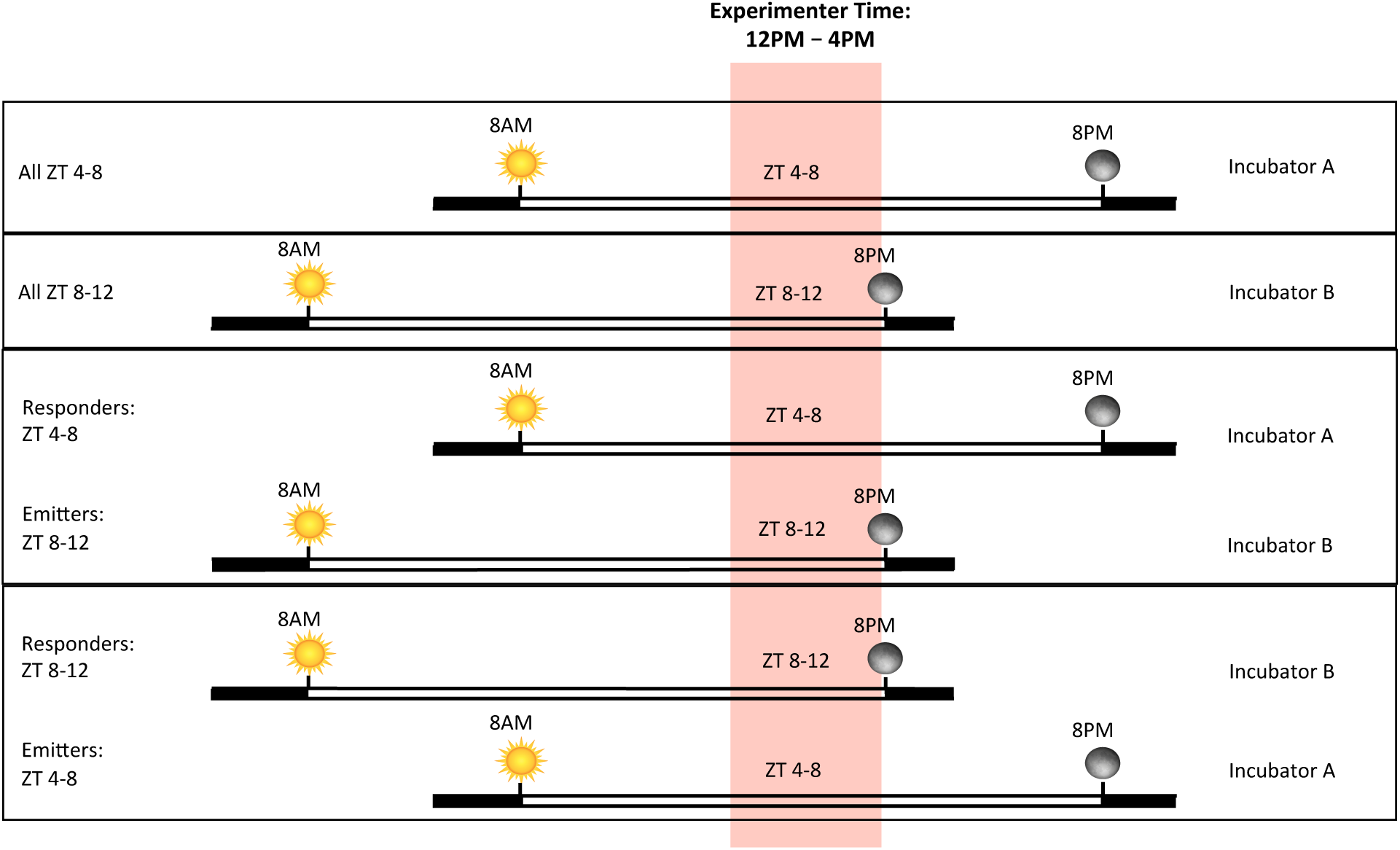
The experimental design used to determine the potential effect of time of day on both the emission and reception of dSO by *D. melanogaster.*

While Fernandez *et al*. (2014) showed that as few as 10 flies can be used as a source of dSO, it is unknown if lower densities produce enough to elicit a response. Therefore, we examined the behaviour of 30 responders when tested with volatile cues produced by 1 (either male or female – data pooled as not statistically different, data not shown), 2, 5 or 10 flies of mixed sex individuals with at least 11 replicates for each emitter age.

We tested the importance of stress intensity comparing the response of 15 mixed sex responders to dSO from 20 emitters that had been submitted to either 1, 2 or 3 bouts (15 sec) of vortexing, in 1 min or that had been transferred to the vial by shaking. We also compared the response of both males and females to the odour from 20 flies (10 males/10 females) that had not been stressed (transferred using the response to light as described above), vortexed for 1 min or held in a vial without agitation or food for 12h. In all cases there were at least 9 replicates per treatment.

To test the dissipation of dSO we compared the response of both males and females to sources immediately following, or 1 or 2 min after the emitters had been vortexed and removed. We also examined the time after vortexing that dSO was emitted. In this case the flies were vortexed, allowed to rest for 10 sec, 1 min, 1 or 2h before being transferred to a clean vial for 1 min. The flies were introduced and removed from the test vial via the light response to ensure there was no additional agitation. In all assays there were 9 replicates.

As there is evidence that the dSO emitted by *D. melanogaster* contains components other than CO_2_, (Suh et al., 2004), we conducted assays comparing the intra and interspecific responses of *D. melanogaster, D. simulans* and *D. suzukii*, with a minimum of 9 replicates for each combination.

### Statistical Analysis

One-way and Two-way ANOVAs were used, followed by Tukey’s *post hoc* test to correct for multiple comparisons in GraphPad Prism (version 7.0a for Mac, GraphPad Software, La Jolla California USA, www.graphpad.com). In all assays with an air vials as a control, we used Welch’s t-tests to confirm that the response to air was not significantly different from 0, and as this was always the case the values were not included. A Welch’s t-test was also used to compare the effect of aged males on the response to dSO.

## RESULTS

The response was not affected by either sex (*F*_1,32_=0.9248, *P*=0.3434) or mating status *F*_1,32_=4.098, *P*=0.0514) when 30 responders were tested to the dSO from 70 emitters (**Figure 2A,B).**

**Figure 2.**
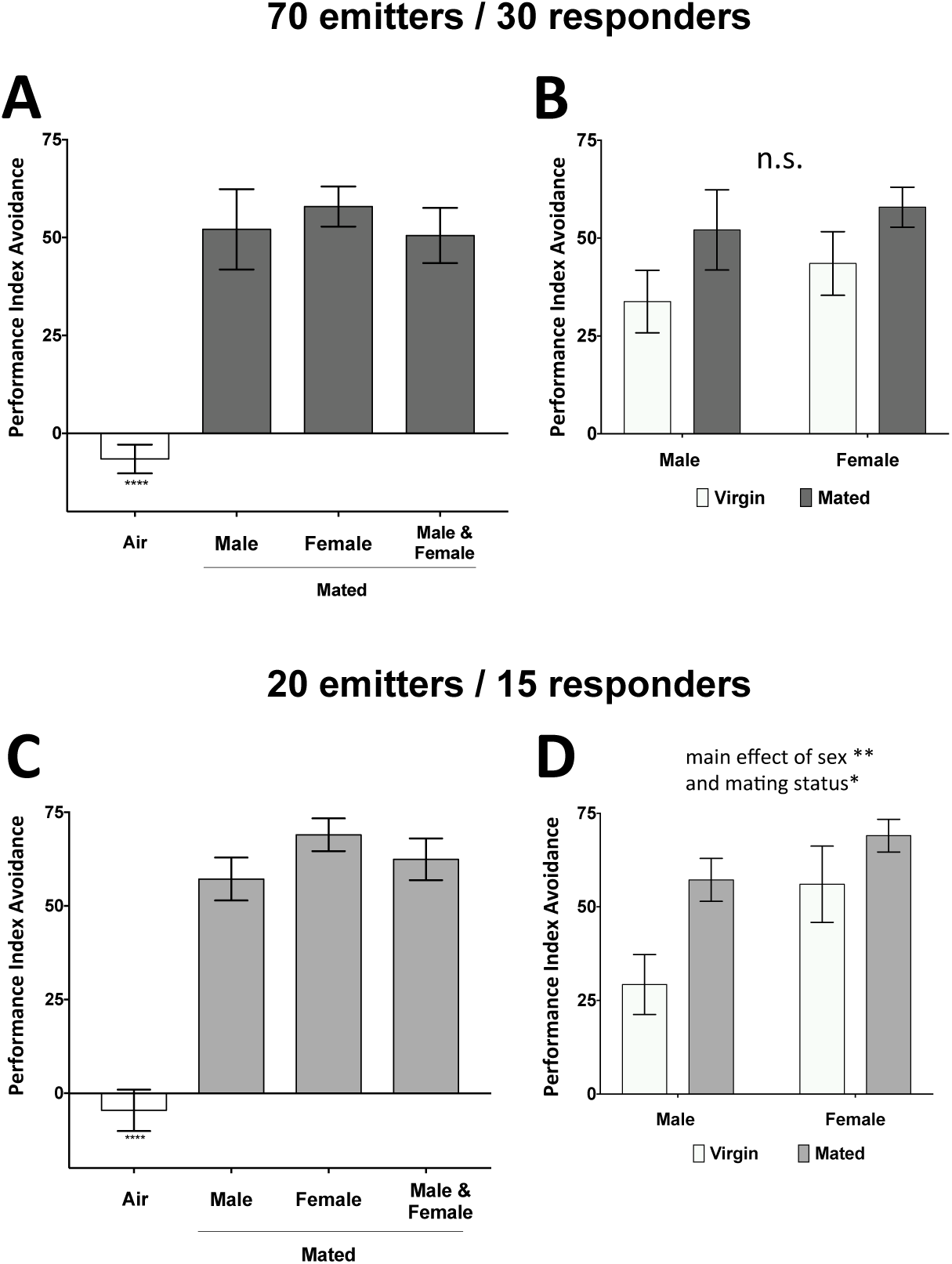
The effect of sex (A and C) and mating status (B and D) on the response of *D. melanogaster* adults to dSO at two different densities of emitters and responders.

However, the response of 15 responders to the dSO from 20 emitters for both sex (*F*_1,29_=7.191, *P*=0.0120) and mating status (*F*_1,29_=8.088, *P*=0.0081) were significant. Females showed a greater response than males (that was also marginally the case at the higher density) and in both sexes mated individuals were more responsive than virgins (**Figure 2C, D**).

While all response levels differed significantly from the air control, post hoc tests showed the age of emitters did not affect the response of responders (**Figure 3A**; *F*_8,81_=42.64, *P*<0.0001), Similarly, responders responded at the same level regardless of the sex of the emitters (**Figure 3B**, *F*_1,28_=0.01086, *P*=0.9177), and there were no differences observed in males and female responders (Figure 3B; *F*_1,28_=1.913, *P*=0.1776). However, when comparing young (3-4 days) versus older (7-10 old) responders there was overall effect of sex with older males showing a significantly lower response that younger ones (**Figure 3C**; *t*_*14.19*_ =2.434, *P*=0.0287).

**Figure 3.**
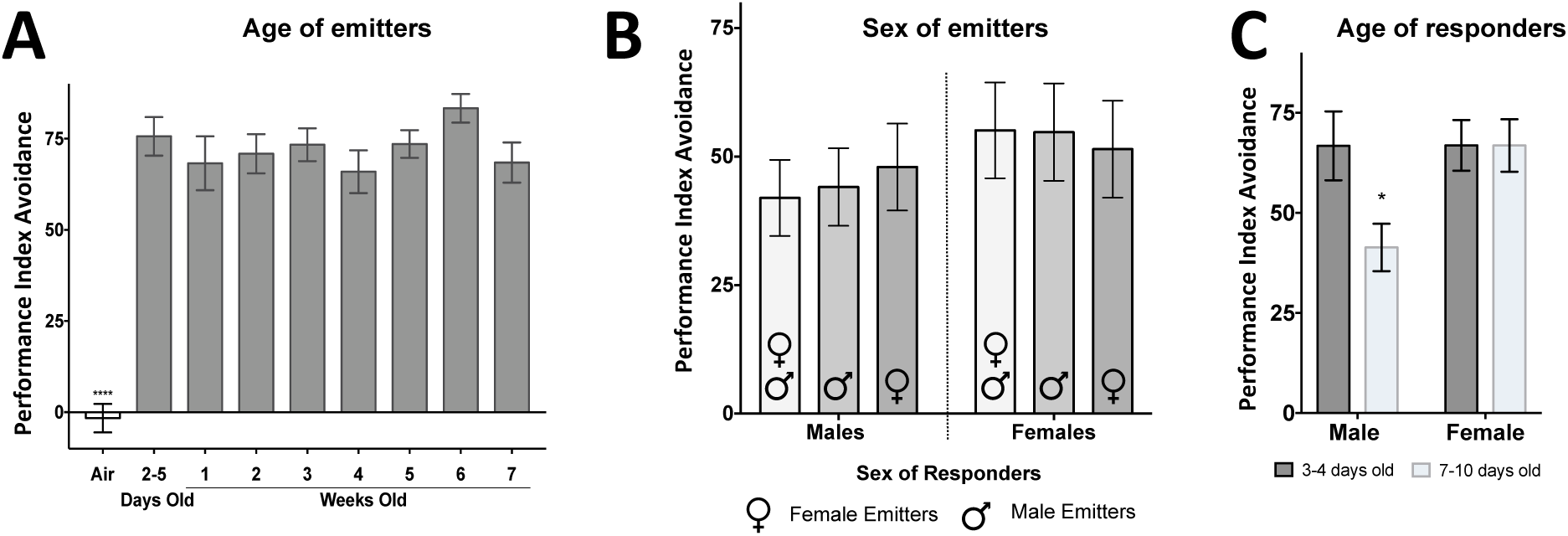
The effect of age (A) and sex (B) of emitters, as well as the age of responders (C) on the response of *D. melanogaster* adults to dSO.

There was no difference in the levels of response when testing both emitters and responders at different times during their photophase (**Figure 4**; *F*_3,64_=0.3224, *P*<0.8091), however, females showed a significantly higher response that males (**Figure 4**; *F*_1,64_=8.084, *P*<0.006).

**Figure 4.**
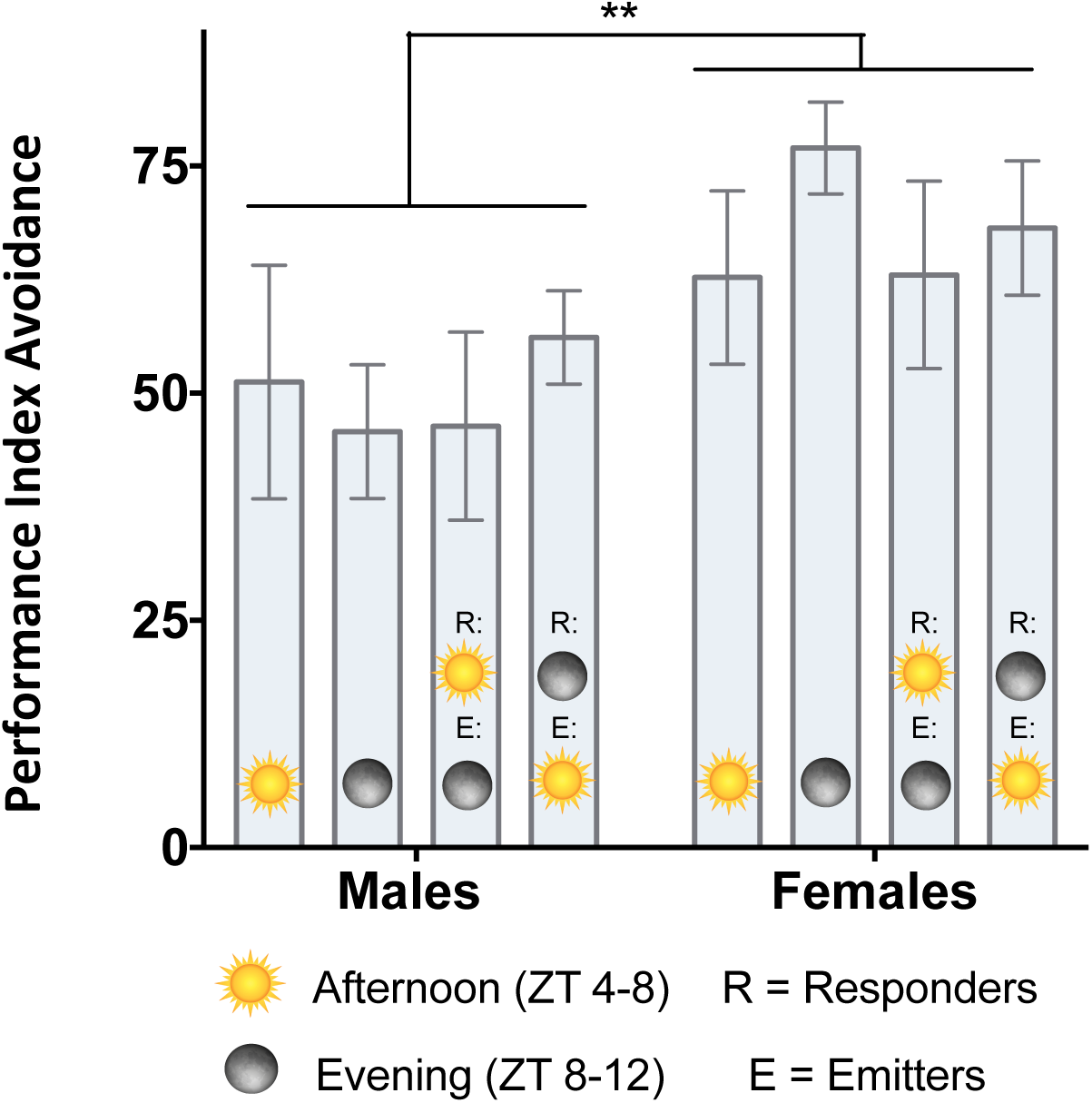
The effect of the time during the photophase that emitters and responders are tested on the response of *D. melanogaster* adults to dSO.

While all dSO sources differ significantly from the air control, the level of response observed is not significantly affected by the number of flies used to generate the odour cue (Figure 5A; F4,92=4.2, P=0.0038). Just the mechanical transfer of flies from one vial to another results in the same level of avoidance as vortexing emitters one to three times in a minute, all of which are significantly higher than the air control (**Figure 5B**; *F*_4,45_=29.45, *P*<0.0001). This is supported by the fact that the response to unstressed flies (transferred using their response to light) was significantly less than to disturbed flies and those stressed by being held without food/water for 12h (**Figure 5 C**; *F*_3,118_ = 101.8, *P*<0.0001) Again, females showed a significantly higher response than males (**Figure 5C**; *F*_*1,118*_ = 7.257, *P=*0.008).

**Figure 5.**
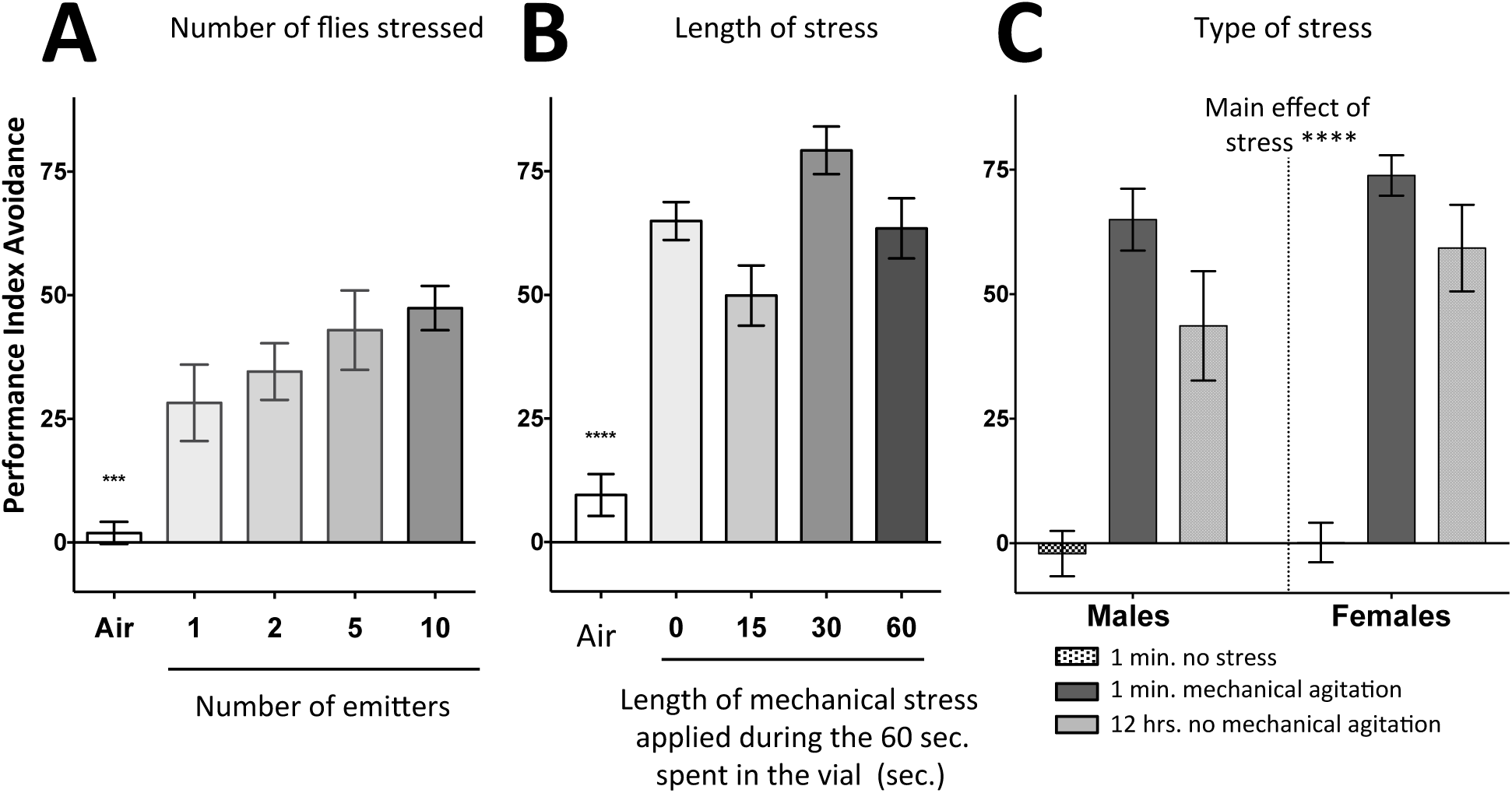
The effect of density and the form of stress emitters were subject to on the response of *D. melanogaster* adults to dSO.

The time elapsed between generating the dSO source and testing it significantly affected the level of avoidance behaviour observed (**Figure 6A**; *F*_*2,60*_=41.43, *P<0.0001*). Both males and females showed a significant level of avoidance when exposed to a dSO source 1 min after it was generated but not after 2 min, indicating the olfactory cue has a high level of volatility. However, the time flies emitted dSO following agitation varied (**Figure 6**B; *F*_4,80_=36.14, *P*<0.0001), with both males and females emitting dSO for at least 1h, but after 2h responses elicited were not different than the air controls. As in other experiments, females showed a significantly higher avoidance than males (*F*_1,80_=6.446, *P*<0.0001).

**Figure 6.**
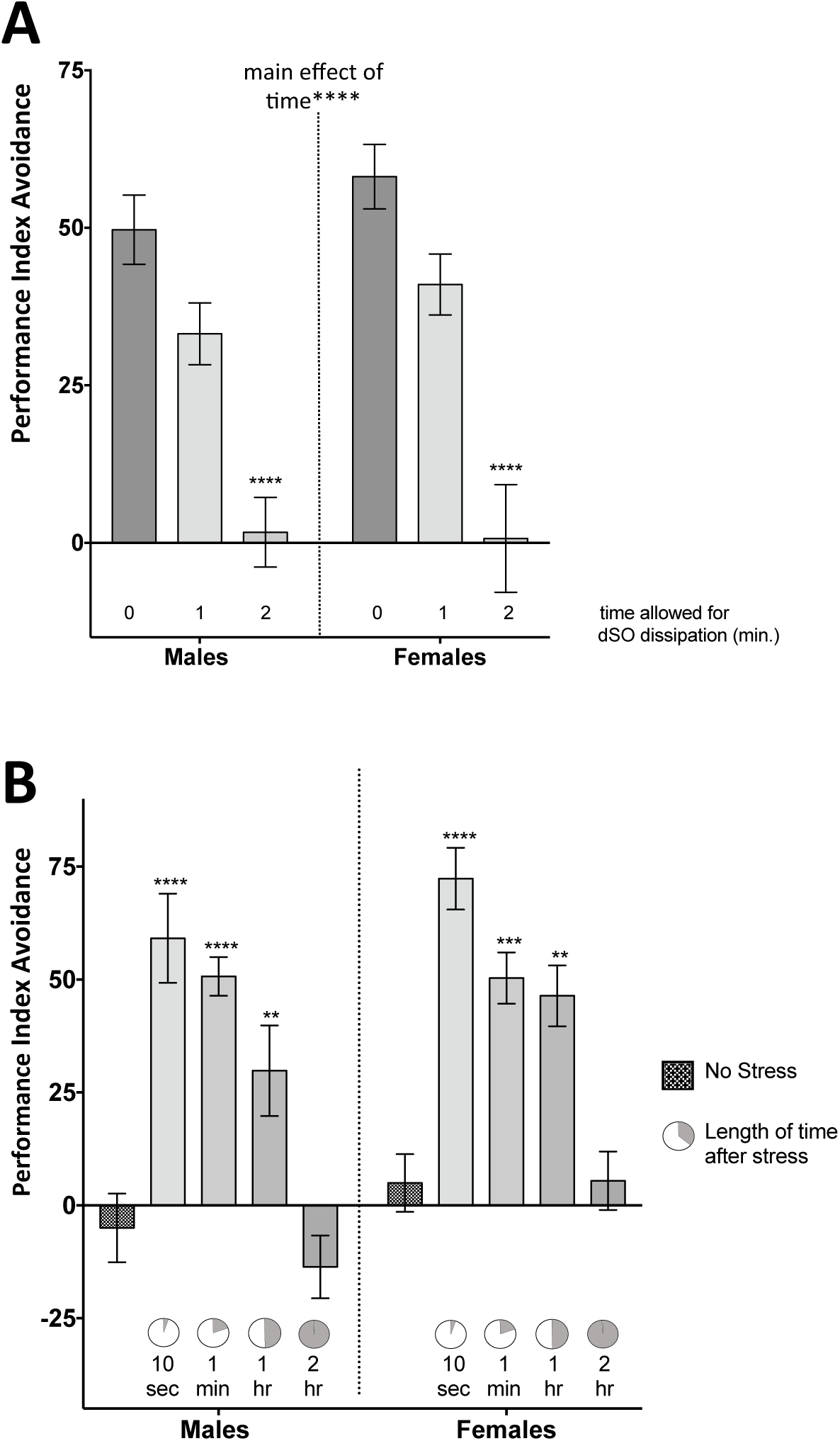
The effect of time on the persistence of dSO (A) and the time since emitters were agitated on the response of *D. melanogaster* adults to dSO.

All three species exhibited some level of response to odours from conspecific and heterospecific sources but there were significant interspecific differences (**Figure 7**; *F*_8,71_=10.5, *P*<0.0001). For example, *D. melanogaster* showed strong responses to all three sources, while *D. susukii* exhibited lower responses to all three. In both species, the level of response did not differ between intra or interspecific sources (*D. melanogaster F*_2,25_=1.339, *P*=0.2803; *D. suzukii F*_2,25_=0.5949, *P*=0.5593). Interestingly, *D. simulans* was the only species that showed the highest response to the conspecifics odour source, which was marginally significant (*F*_2,21_= 3.277, *P*=0.0577).

**Figure 7.**
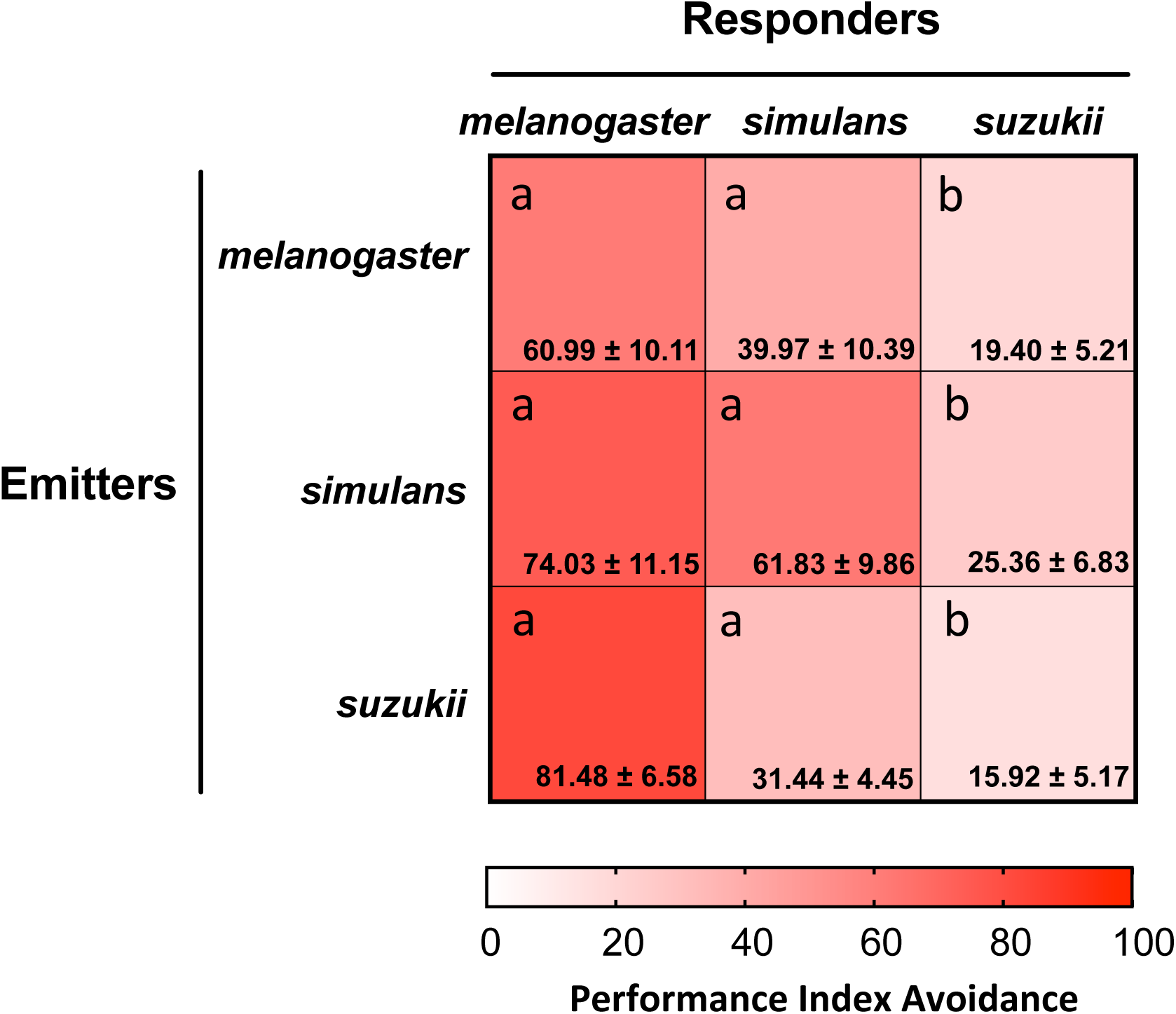
A heat map comparing the response of *D. melanogaster, D. simulans* and *D.susukii* adults to the dSO emitted by conspecific and heterospecific adults. The analysis compared all values with the *D. melanogaster* response to conspecifics.

## DISCUSSION

Our results support the idea that *D. melanogaster* adults emit olfactory cues that may affect the behaviour of conspecifics whenever they are subjected to any form of stress. In part this is probably the result of physiological changes, as seen in bees that had been vortexed (Bateson, Desire, Gartside, & Wright, 2011), that increases the production of CO_2_ which is a major component of dSO (Suh et al., 2004).

However, the response to alarm signals is contextual, probably because of trade-offs between the associated costs and benefits related to reproductive success, as clearly shown in the responses of different stages of the green peach aphid to (E)-β-farnesene (Montgomery & Nault, 1978). This is also the case for *D. melanogaster* responding to dSO. For example, the lower responses by virgin flies compared to mated individuals may relate to mating opportunities, given that the estimated mean longevity of *Drosophila* adults under field conditions is about 3-7 days (Rosewell & Shorrocks, 1987). Thus, while both virgin and mated individuals leaving a site would benefit from avoiding potential danger, virgins leaving a site where there are conspecifics decrease their chances of acquiring a mate.

The data from several of our experiments show that mated males exhibit lower response levels than mated females of the same age, which again could relate to aspects of reproductive success. The more mates a male acquires the higher his potential lifetime reproductive output (e.g. Royer & McNeil, 1993), thus a lower response to dSO may increase his chances of encountering a receptive female. Similarly, being in a dangerous site may reduce short term oviposition opportunities for a mated female, given that one mating provides her enough sperm to fertilize her full egg compliment, leaving would increase her chances of surviving and increasing opportunities for future oviposition. Similarly, the absence of an age-related response to dSO in females is not surprising for although the highest daily egg output occurs during the first week (Tatar, Promislow, Khazaeli, & Curtsinger, 1996), moving away from potential danger could extend her future opportunities to oviposit. In contrast, as mating opportunities for males decline with age (see Ruhmann, Koppik, Wolfner, & Fricke, 2018) older males would have less to lose than younger ones with respect to future reproduction.

The responses by *D. melanogaster* and *D. simulans* to both conspecific and heterospecific cues were always high, while *D. suzukii* has low responses to all sources. Krause Pham and Ray (2015) noted that younger fruits emit higher levels of CO_2_ and postulated the low responsiveness of *D. suzukii* adults would facilitate foraging for suitable sites. In addition, the high abundance and the distribution of available feeding/oviposition sites under natural ecological conditions would normally result in low spatial densities of *D. suzukii*, so the relative benefit gained by responding to a rapidly dissipating conspecific alarm cue would be of limited value. In contrast, the relative densities of *D. melanogaster* and *D. simulans* would be higher on fermenting fruits so there would be a higher probability of detecting dSO from conspecifics in proximity and responding to a danger source that is close by would be advantageous.

The only species that showed a higher response to conspecific rather than heterospecific ones, was *D. simulans*, something one would expect if, in addition to CO_2_, there are species specific components in dSO (Enjin & Suh, 2013; Suh et al., 2004). Even if there is species specificity, the high responses of *D. melanogaster* and *D. simulans* to heterospecifics suggests that there is enough CO_2_ alone emitted by stressed flies to elicit responses and/or that the two species share other dSO common components. Clearly, additional research is required to identify if there are other components in the dSO from each species, and to determine to what extent they alter avoidance behaviours. Furthermore, once all components have been identified, it will be possible to determine to what extent profiles change with age, mating status or the type of stress that the files are subjected to. This information will be important when investigating the potential use of dSO in pest management programmes against species such as *D. melanogaster*, although our results suggest such an approach would not be as effective against *D. suzukii.*

*D. melanogaster* are a great model for neuro-genetic studies due to the availability of insertion and deletions that cover most of the fully identified genome, RNAi libraries, and transgenic lines to repress or enhance the expression of certain genes (Hales, Korey, Larracuente, & Roberts, 2015). Our results provide a better understanding of the parameters that should be considered in future work looking at the dSO detection, research that could lead to a better general understanding of neurophysiological aspects of insect olfaction.

The findings also show that the regular techniques used to transfer flies will result in the release of dSO and that while the olfactory cue dissipates quickly, stressed flies may continue to emit for at least an hour. These effects of manipulation could be important confounding factor when studying certain aspects of behaviour, as seen in studies examining aggression (Trannoy, Chowdhury, & Kravitz, 2015). Consequently, we would suggest that anyone conducting research on *Drosophila* behaviour use the light response approach when transferring flies.

## ACKNOWLEDGEMENTS

This work was supported by a Western Science undergraduate pre-thesis award and an internal graduate scholarship to RTY, Western Foundation internal grant and NSERC Discovery Grants 04507-2015 to JNM and 04275-2015 to AFS. We thank Ian S. McDonald for his technical contribution, Yanira Jimenez Padilla and the Sinclair lab for providing the food for *Drosophila suzukii* during the length of the experiment.

## AUTHOR CONTRIBUTIONS

Data acquisition and analysis were carried out by RTY, EL, ISM, SC, ISM, AFG and MS; Protocols elaborated by AFS and JNM; the first draft was written by RTY, EL and AFS and the final by RTY, AFS and JNM.

## ETHICAL NOTE

No approval is required from the Western’s Animal Care Committee or the Provincial and Federal regulatory bodies to study invertebrates. However, we provided appropriate rearing conditions and anesthetized flies using CO_2_ or cold anaesthesia when manipulating flies for colony maintenance. Stress treatments to produce dSO resulted in no mortality

## REFERENCES

Bateson, M., Desire, S., Gartside, S. E., & Wright, G. A. (2011). Agitated Honeybees Exhibit Pessimistic Cognitive Biases. Current Biology, 21(12), 1070–1073.

Benzer, S. (1967). Behavioral mutants of *Drosophila melanogaster* isolated by countercurrent distribution. PNAS, 58, 1112–1119.

Bracker, L. B., Siju, K. P., Varela, N., Aso, Y., Zhang, M., Hein, I.,… Grunwald Kadow, I. C. (2013). Essential role of the mushroom body in context-dependent CO_2_ avoidance in Drosophila. Curr Biol, 23(13), 1228–1234. doi:10.1016/j.cub.2013.05.029

Brenman-Suttner, D. B., Long, S. Q., Kamisan, V., de Belle, J. N., Yost, R. T., Kanippayoor, R. L., & Simon, A. F. (2018). Progeny of old parents have increased social space in *Drosophila melanogaster*. Scientific Reports(8), 3673. doi:10.1038/s41598-018-21731-0

Chao, Y. C., Fleischer, J., & Yang, R. B. (2018). Guanylyl cyclase-G is an alarm pheromone receptor in mice. EMBO J, 37(1), 39–49. doi:10.15252/embj.201797155

Dahanukar, A., & Ray, A. (2011). Courtship, aggression and avoidance: Pheromones, receptors and neurons for social behaviors in Drosophila. Fly, 5(1), 58–63. doi:10.4161/ y.5.1.13794

Dubruille, R., & Emery, P. (2008). A plastic clock: how circadian rhythms respond to environmental cues in Drosophila. Mol Neurobiol, 38(2), 129–145. doi:10.1007/s12035-008-8035-y

Enjin, A., & Suh, G. S. (2013). Neural mechanisms of alarm pheromone signaling. Mol Cells, 35(3), 177–181. doi:10.1007/s10059-013-0056-3

Faucher, C., Forstreuter, M., Hilker, M., & de Bruyne, M. (2006). Behavioral responses of Drosophila to biogenic levels of carbon dioxide depend on life-stage, sex and olfactory context. J Exp Biol, 209(Pt 14), 2739–2748. doi:10.1242/jeb.02297

Fernandez, R. W., Akinleye, A. A., Nurilov, M., Feliciano, O., McDonald, I. S., & Simon, A. F. (2014). Straightforward Assay for Quantification of Social Avoidance in *Drosophila melanogaster*. J. Vis. Exp., 94, e52011. doi:10.3791/52011

Hales, K. G., Korey, C. A., Larracuente, A. M., & Roberts, D. M. (2015). Genetics on the fy: A primer on the Drosophila model system. Genetics, 201(3), 815–842. doi:10.1534/genetics.115.183392

Hunt, G. J. (2007). Flight and fight: a comparative view of the neurophysiology and genetics of honey bee defensive behavior. J Insect Physiol, 53(5), 399–410. doi:10.1016/j.jinsphys.2007.01.010

Jakobs, R., Ahmadi, B., Houben, S., Gariepy, T. D., & Sinclair, B. J. (2017). Cold tolerance of third-instar *Drosophila suzukii* larvae. Journal of Insect Physiology, 96, 45–52. doi:https://doi.org/10.1016/j.jinsphys.2016.10.008

Krause Pham, C., & Ray, A. (2015). Conservation of olfactory avoidance in Drosophila species and identification of repellents for *Drosophila suzukii*. Sci Rep, 5, 11527. doi:10.1038/srep11527

Kwon, J. Y., Dahanukar, A., Weiss, L. A., & Carlson, J. R. (2007). The molecular basis of CO2 reception in Drosophila. Proceedings of the National Academy of Sciences U S A, 104(9), 3574–3578. Epub 2007 Feb 3520.

Mathuru, A. S., Kibat, C., Cheong, W. F., Shui, G., Wenk, M. R., Friedrich, R. W., & Jesuthasan, S. (2012). Chondroitin fragments are odorants that trigger fear behavior in fish. Curr Biol, 22(6), 538–544. doi:10.1016/j.cub.2012.01.061

Montgomery, M. E., & Nault, L. R. (1978). Effects of age and wing polymorphism on the sensitivity of *Myzus persicae* to alarm pheromone. Annals of the Entomological Society of America, 71(5), 788–790. doi:10.1093/aesa/71.5.788

Mujica-Parodi, L. R., Strey, H. H., Frederick, B., Savoy, R., Cox, D., Botanov, Y.,… Weber, J. (2009). Chemosensory cues to conspecific emotional stress activate amygdala in humans. PLoS One, 4(7), e6415. doi:10.1371/journal.pone.0006415

Napper, E., & Pickett, J. A. (2008). Alarm Pheromones of Insects. In J. L. Capinera (Ed.), Encyclopedia of Entomology (pp. 85–95). Dordrecht: Springer Netherlands.

Rosewell, J., & Shorrocks, B. (1987). The implication of survival rates in natural populations of Drosophila: capture-recapture experiments on domestic species. Biological Journal of the Linnean Society, 32, 373–384.

Royer, L., & McNeil, J. N. (1993). Male Investment in the European Corn Borer, *Ostrinia nubilalis* (Lepidoptera: Pyralidae): Impact on Female Longevity and Reproductive Performance. Functional Ecology, 7(2), 209–215. doi:10.2307/2389889

Ruhmann, H., Koppik, M., Wolfner, M. F., & Fricke, C. (2018). The impact of ageing on male reproductive success in *Drosophila melanogaster*. Exp Gerontol, 103, 1–10. doi:10.1016/j.exger.2017.12.013

Siju, K. P., Bracker, L. B., & Grunwald Kadow, I. C. (2014). Neural mechanisms of context-dependent processing of CO_2_ avoidance behavior in fruit flies. Fly (Austin), 8(2), 68–74. doi:10.4161/fly.28000

Sokolowski, M. B. (2010). Social interactions in “simple” model systems. Neuron, 65(6), 780–794.

Suh, G. S., Ben-Tabou de Leon, S., Tanimoto, H., Fiala, A., Benzer, S., & Anderson, D. J. (2007). Light activation of an innate olfactory avoidance response in Drosophila. Curr Biol, 17(10), 905–908. doi:10.1016/j.cub.2007.04.046

Suh, G. S., Wong, A. M., Hergarden, A. C., Wang, J. W., Simon, A. F., Benzer, S.,… Anderson, D. J. (2004). A single population of olfactory sensory neurons mediates an innate avoidance behaviour in Drosophila. Nature, 431(7010), 854–859. doi:10.1038/nature02980

Tatar, M., Promislow, D. E. L., Khazaeli, A. A., & Curtsinger, J. W. (1996). Age-Specific Patterns of Genetic Variance in *Drosophila melanogaster.* II. Fecundity and Its Genetic Covariance With Age-Specific Mortality. Genetics, 143(2), 849–858.

Trannoy, S., Chowdhury, B., & Kravitz, E. A. (2015). Handling alters aggression and “loser” effect formation in *Drosophila melanogaster*. Learn Mem, 22(2), 64–68. doi:10.1101/lm.036418.114

Turner, S. L., & Ray, A. (2009). Modification of CO_2_ avoidance behaviour in Drosophila by inhibitory odorants. Nature, 461(7261), 277–281. doi:10.1038/nature08295

Vandermoten, S., Mescher, M. C., Francis, F., Haubruge, E., & Verheggen, F. J. (2012). Aphid alarm pheromone: an overview of current knowledge on biosynthesis and functions. Insect Biochem Mol Biol, 42(3), 155–163. doi:10.1016/j.ibmb.2011.11.008

Verheggen, F. J., Haubruge, E., & Mescher, M. C. (2010). Alarm Pheromones—Chemical Signaling in Response to Danger. In G. Litwack (Ed.), Vitamins & Hormones (Vol. 83, pp. 215–239): Academic Press.

Yew, J. Y., & Chung, H. (2015). Insect pheromones: An overview of function, form, and discovery. Prog Lipid Res, 59, 88–105. doi:10.1016/j.plipres.2015.06.001

Zhou, Y., Loeza-Cabrera, M., Liu, Z., Aleman-Meza, B., Nguyen, J. K., Jung, S. K.,… Zhong, W. (2017). Potential nematode alarm pheromone induces acute avoidance in *Caenorhabditis elegans*. Genetics, 206(3), 1469–1478. doi:10.1534/genetics.116.197293

